# Selective facilitation of egocentric mental transformations under short-term microgravity

**DOI:** 10.1101/851162

**Authors:** Nicolas Meirhaeghe, Virginie Bayet, Pierre-Vincent Paubel, Claudine Mélan

**Affiliations:** Harvard-MIT Division of Health Sciences & Technology, Cambridge, Massachusetts, USA; Laboratoire CLLE, CNRS 5263, University Toulouse II Jean Jaurès, Toulouse, France

**Author notes:** **Correspondence**, Nicolas Meirhaeghe, Harvard-MIT Division of Health Sciences & Technology, 43 Vassar Street, Building 46-6041, Cambridge, MA 02139, USA. **Author contributions** C.M. and N.M. conceived the project. N.M. collected pilot data and performed preliminary analyses used to refine the task design. P-V.P. and N.M. adapted the experiment to virtual reality. P-V.P. and C.M. handled technical aspects of the parabolic flights. All authors took part in the parabolic flights and contributed equally to data collection. P-V.P. and N.M. processed the parabolic flight data. N.M., V.B., and C.M. performed the statistical analyses. All authors were involved in interpreting the results. N.M. and C.M. wrote the manuscript. **Declaration of interests** The authors declare no competing interests.

**Keywords:** Egocentric transformation, mental rotation, perspective-taking, microgravity, parabolic flight, virtual reality

## Abstract

Understanding the impact of microgravity on human cognitive performance is crucial to guarantee the safety and success of future long-term manned missions. The effects of weightlessness on key mental processes such as spatial abilities are in particular not fully characterized. In this study, we examine the influence of microgravity on perspective-taking abilities—a type of mental operation especially relevant in the context of collaborative teamwork between ‘free-floating’ astronauts. Twelve subjects performed a cooperative task in virtual-reality under both normal and short-term microgravity conditions during a parabolic flight. The task involved various degrees of mental transformations, and required subjects to perform actions instructed by a fellow astronaut aboard a virtual spacecraft. The experimental design allowed us to control for nuisance variables, training effects, and non-gravity related factors of parabolic flights. Overall, our results indicated that microgravity has a facilitatory effect on perspective-taking abilities. Notably, this facilitation was selective to conditions requiring subjects to rotate their perspective around their line of sight, i.e., for mental rotations in the frontal plane. Moreover, microgravity affected subjects differently depending on their visual field dependence, as determined via a classic rod-and-frame test. Specifically, improvement in performance was more pronounced in field-independent subjects. Together, these results shed light on a long standing debate about the impact of microgravity on egocentric mental imagery, and have direct operational consequences for future long-term missions.

## 1. INTRODUCTION

Gravity plays an integral part in countless aspects of human behavior. When interpreting complex visual scenes, interacting with novel objects, walking around or simply reasoning about the physical world around us, we implicitly rely on gravitational cues to guide our decisions and actions. Gravity can be considered as a force acting both on the vestibular and proprioceptive systems, providing a constant reference to which we anchor our mental representations of the environment [1]. For instance, the perception of ‘upright’ is the result of integrating the direction of perceived gravity with an internal representation of the body’s longitudinal axis and the orientation of the ambient visual world [2, 3]. Depriving individuals from a gravitational reference, as in space or parabolic flights, is therefore expected to alter their ability to function and rely on these mental representations [4]. Prior studies have indeed reported a wide range of debilitating effects in weightlessness affecting both sensorimotor [5–8] and cognitive functions [9, 10]. Crucial abilities such as depth perception [11, 12], visual recognition [13], motor imagery [14], estimation of body orientation [15] and mental rotation [16] have all been shown to be impaired. These deficits have raised serious concerns for future long-term manned missions, and have underlined the need for a better understanding and characterization of human performance under both short and long-term microgravity exposure [13, 17].

During space missions, astronauts perform teleoperation tasks and cooperate with one another by relying on a mental representation of their surroundings. For instance, they may need to perform mental rotations to identify a desired tool floating amongst other objects, or they may need to mentally adopt a different perspective to visualize the equipment installed in a different cabin or assess the orientation of an instrument being remotely controlled. Furthermore, rather than simply adopting another spatial point of view (called primary or 1^st^-person perspective; [18]), astronauts may need to mentally adopt a team mate’s point of view while performing a collaborative task. In this situation, a social component is added to the mental transformation, as it now relies on an interpretation of the other person’s intentions [19], and is referred to as a 3^rd^-person perspective. Astronauts may encounter difficulties performing these types of transformation [20] when confronted with conflicting visual and vestibular information. In particular, the up-down orientation indicated by cabin interiors providing visual landmarks (allocentric reference) may differ from the one determined by the astronauts’ longitudinal body-axis (egocentric reference) while they are floating in the cabin. Given how ubiquitous mental rotation and perspective-taking situations are in space operations, it appears essential to investigate the effects of weightlessness on these processes, particularly when visual landmarks and body-orientations differ.

Although the impact of weightlessness on spatial abilities has been extensively investigated, studies on the specific case of mental rotation and perspective-taking remain relatively limited (see [16] for a review). Early studies conducted on astronauts found that performance in mental rotation tasks was facilitated in microgravity [21–23]. Notably, improvements appeared more pronounced for rotations in the frontal plane, that is, when subjects were imagining objects rotating around their line of sight [22]. From a theoretical standpoint, the impact of microgravity on mental rotations were accounted for by the concept of motoric embodiment [24], which postulates that spatial abilities involve a mental simulation of the actual movements necessary to perform a desired transformation [25, 26]. This view may explain why mental rotations in the frontal plane are facilitated in microgravity. Because these rotations correspond precisely to movements that are nearly impossible on Earth, but become conceivable in a weightless environment, the release of physical constraints on the mental simulation process in 0g may lead to a performance enhancement. In line with this interpretation, it has been shown that a complex visual background presented in different orientations affected the perceptual vertical less in microgravity than upright in normal gravity [27].

Despite the theoretical and experimental support for a positive influence of microgravity on mental rotation abilities, conclusions have nevertheless remained highly nuanced. Later studies indeed contradicted the original results and found no evidence for an impact of weightlessness on mental rotations [28–30], while others actually found a decline in performance [16, 31]. The reasons behind these conflicting findings are not entirely clear. One possibility is that they partly stem from variations in the experimental setup used to test mental rotation abilities [31]. Alternatively, the observed effects—or absence thereof—could also arise from non-specific factors characteristic of weightless environments such as stress, high workload, lack of sleep and physiological changes, that are not directly related to gravity [32]. In fact, it was previously suggested that facilitation or impairment in cognitive performance during weightlessness phases of parabolic flights compared to ground observations may be attributable to increased stress rather than weightlessness itself [13]. In that regard, [33] argued that in-flight stress effects could be controlled for by performing measurements on repetitive and consecutive periods of normal gravity and microgravity.

Yet another explanation for the contradicting findings relates to differences in strategy that subjects may have used to solve the tasks. Indeed, depending on the reference frames being manipulated, i.e, if subjects imagine themselves moving relative to the environment (egocentric) or imagine instead objects in the environment moving with respect to them (allocentric), performance characteristics including response times can differ [34, 35]. Moreover, there is evidence that egocentric transformations are more impacted by changes in vestibular inputs [36, 37], including those induced by microgravity [29], compared to allocentric transformations. Based on these observations, it was argued that the presence of a gravitational frame of reference might facilitate egocentric transformations [16]. Finally, the propensity of subjects to rely on or be distracted by visual cues during mental tasks [38] is known to vary across individuals [39]. This visual field dependency may modulate subjects’ performance, especially in tasks requiring the processing of visual versus proprioceptive and vestibular information.

Besides controversial findings, methodological limitations have also contributed to opacifying the question of mental imagery in weightlessness. Prior studies have indeed been limited to simplistic stimuli, including abstract 3D objects [22, 28] used in classic spatial tasks [40], drawings of whole body [31, 41], body parts [31], or faces [42]. Although these stimuli offered greater control over behavioral variables, the artificial nature of the tasks employed has precluded a full characterization of the impact of microgravity in a more realistic, operationally-relevant context. In particular, more complex mental transformations such as perspective-taking abilities involving more than one agent remain fully uncharacterized. This consideration is especially relevant in the prospect of future long-term missions likely to embark large teams of individuals who will have to coordinate complex tasks [17].

In this study, we sought to examine the impact of microgravity on perspective-taking abilities. The main goal was to determine whether these transformations are degraded, facilitated, or unaffected in weightlessness. In contrast with prior studies, we addressed this question from an operational standpoint by considering a realistic paradigm meant to reproduce typical interactions among free-floating astronauts. Accordingly, we developed a novel experiment in virtual reality that required trained subjects to perform a complex cooperative task. The task crucially depended on subjects’ ability to mentally project themselves in different locations and orientations inside a virtual spacecraft. Participants completed the task aboard a parabolic flight aircraft both in normal gravity conditions and under short-term microgravity. According to the motoric embodiment hypothesis, performance at the perspective-taking task was expected to *improve* in weightlessness, especially for rotations within the frontal plane (i.e., roll rotations), as a consequence of the release of physical constraints on the mental rotation processes. Alternatively, if perspective-taking normally benefits from the presence of a gravitational reference frame, performance was expected to *drop* in microgravity. The task included several control conditions allowing us to disentangle the impact of microgravity purely on the integration of visual landmarks and to control for other nonspecific factors inevitably introduced in parabolic flight studies. Finally, we also tested whether subjects’ dependence on the visual field (i.e., their tendency to rely on visual cues to assess verticality) modulated the potential effects of microgravity on performance.

## 2. METHOD

### 2.1. Subjects

Twelve participants (all males, aged 35–49) took part in the study. All participants had normal or corrected-to-normal vision and reported no history of vestibular dysfunction. All had given their informed consent in accordance with the ethical standards of the Declaration of Helsinki. Subjects had passed medical tests to qualify for the parabolic flights (i.e., equivalent of an Air Force Class III medical examination). All subjects had one prior experience of parabolic flights except for one participant who had participated in two prior parabolic flights. They all received a standard dose of scopolamine to prevent motion sickness. One subject became sick during the flight despite medication and was excluded from all the analyses. In addition to the main perspective-taking task, each subject completed a computerized version of the classic rod-and-frame test (RFT; [43]) to assess visual field dependency. The main task and the RFT were implemented in Matlab 2014b (MathWorks, Natick, USA) and ran on a Dell Alienware Cassini 15 computer.

### 2.2. Experimental procedure

The parabolic flight campaign consisted of three flights operated on three successive days. Four participants and two experimenters boarded each flight. Every subject took part in three separate experimental sessions.

*Preflight* session: all subjects completed the RFT test on the first day of the campaign. After familiarization with the test material and the overall procedure, participants performed the main perspective-taking task aboard the aircraft (engine off).

*Inflight* session: on each flight, the aircraft performed a total of 30 parabolas each separated by at least a 2-min period of steady flight (Figure 1d). One parabola consisted of a sequence of 1.8g–0g-1.8g periods, each period lasting 20 seconds. Every five parabolas were followed by a 5-min break in steady flight, extended to 8 min at mid-flight. During each half of the flight, two participants performed the perspective-taking task. For every subject, half the trials were done in microgravity conditions over 20-s periods (inflight 0g). The other half of the trials were done in 1g conditions over 20-s test periods alternating with 20-s rest periods during the 5-min breaks (inflight 1g). This strategy was used to match overall testing duration between 1g and 0g, and to mimic the interruptions between parabolas.

**Figure 1.**
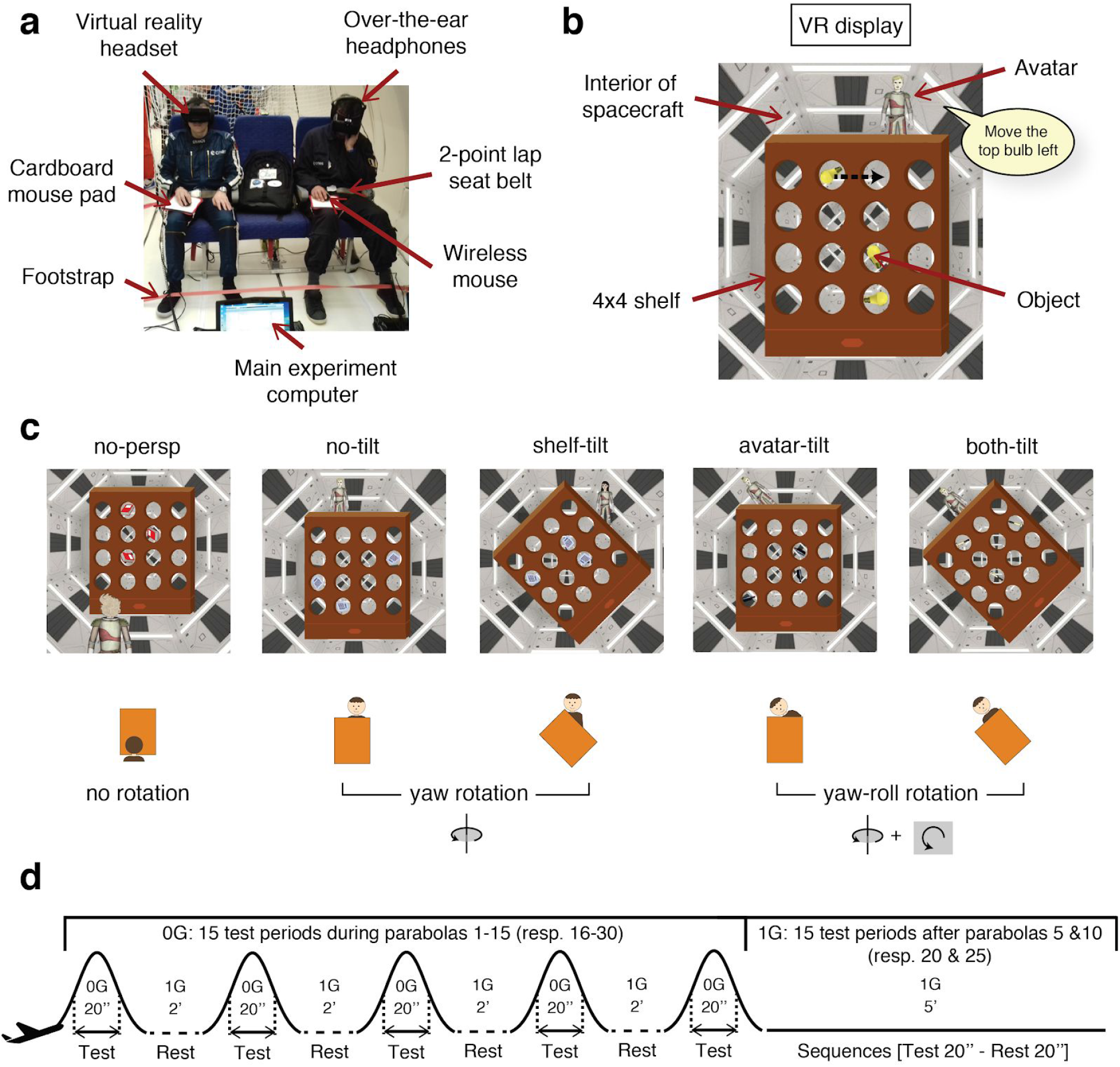
Task design and protocol. a) Typical experimental session aboard the aircraft. Subjects secured in a regular airplane seat were equipped with headphones, a VR headset, and a wireless mouse they manipulated over a solid cardboard directly attached to their thigh. b) Scene shown on the VR display on each trial. Following a brief (1s) appearance, the avatar representing a colleague astronaut instructs the subject through the headphone to move an object from one shelf compartment to another (illustrating by the speech bubble). After interpreting the desired movement, the subject uses the mouse to select the object and place it in the target compartment (dashed black arrow). c) In each of the 5 experimental conditions, the position/orientation of the avatar and/or the orientation of the shelf were manipulated to elicit different degrees of egocentric transformation (no rotation, yaw rotation and yaw-roll rotation). d) Sequence of parabolas for the inflight session. Subjects completed 0g trials (typically 3 or 4) during every 20-s parabola, each separated by a 2-min break. Every 5 parabolas were followed by a longer break (5 min) during which subjects completed 1g trials over alternating 20-s test/rest periods.

*Postflight* session: immediately upon landing, participants performed again the perspective-taking task aboard the aircraft (engine off).

### 2.3. Perspective-taking task

Subjects were seated in a comfortable armchair and secured by footstraps and a 2-point lap seat belt. The straps and seat belt were kept slightly loose over the course of the experiment to reduce tactile and proprioceptive cues during 0g periods. Subjects were equipped with a virtual reality (VR) headset (Oculus Rift DK2), over-the-ear headphones (Sennheiser HD380 PRO), and a wireless mouse that they manipulated over a solid cardboard directly attached to their thigh (Figure 1a). Subjects were instructed to keep their head upright during the whole experiment to ensure that vestibular inputs were constant during 1g periods. In addition, the VR mode was set up such that the image displayed in the headset was stationary with respect to the retina. This guaranteed that the image seen by subjects was completely independent of head motion, which prevented them from solving the perspective-taking task by tilting their head.

Each trial of the experiment started with a fixation cross at the center of the VR display. Following a short delay, a scene depicting the interior of a spacecraft with a 4×4 rectangular shelf floating in the middle appeared on the screen (Figure 1b). The scene covered the entire visual field of the subject to allow maximum VR immersion. On every trial, three identical objects (e.g., three bottles) were placed in individual compartments of the shelf. The location and nature of the objects was randomized on a trial-by-trial basis. After 1 second, an avatar portraying a colleague astronaut popped up either behind or in front of the shelf, in either an upright or tilted (45°) orientation. The exact location of the avatar with respect to the shelf was randomized across trials. The avatar was shown for 1 second before disappearing. Immediately after, a verbal instruction was played in the headphones instructing the participant to move one of the objects from one shelf compartment to another (e.g., “move the top bottle down”). The instruction was always given from the avatar’s perspective. The subject’s goal was to remember the position and orientation of the avatar, and interpret the instructed movement from the appropriate perspective. To respond, subjects had to drag-and-drop the identified object in the desired unit using the mouse. They received no feedback about the accuracy of their responses during the experiment. For each trial, we recorded both the response time (measured from the end of the verbal instruction to the release of the mouse click in the target compartment) and the accuracy of the movement performed (i.e., whether or not the object was moved in the appropriate compartment).

The avatar was only briefly presented (1s) before the instruction to ensure that subjects relied on a mental, as opposed to a purely visual strategy to solve the task. Reports from a pilot study indeed revealed that when the avatar is present throughout the duration of the trial, subjects often use it as a visual reference to determine the left/right and up/down directions. Removing the avatar also ensured that the only visual reference subjects could use was the shelf. The shelf itself was designed to have an inherent up-down polarity (diamond marker at the ‘bottom’ of the shelf), which served as an allocentric visual cue.

The main task was composed of 5 experimental conditions in which either the position/orientation of the avatar, and/or the orientation of the shelf were varied (Figure 1c). In the *no-persp* condition, the avatar appeared upright in front of the shelf, the shelf was also upright. This condition engaged purely visual processes and did not require any mental transformation to be solved. In the *no-tilt* and *shelf-tilt* conditions, the avatar appeared upright behind the shelf, the shelf was respectively upright or tilted. These two conditions required subjects to mentally rotate themselves behind the shelf around the vertical axis (i.e., yaw rotation), regardless of the orientation of the shelf. Finally, in the *avatar-tilt* and *both-tilt* conditions, the avatar appeared tilted behind the shelf, the shelf was respectively upright or tilted in the same direction as the avatar. These two conditions required subjects to mentally rotate themselves around the vertical axis and within the frontal plane behind the shelf (i.e., yaw-roll rotation). The common tilt direction for the avatar and the shelf on the both-tilt condition was balanced across subjects (i.e., 45° clockwise for half the subjects, counterclockwise for the other half). The task originally contained a sixth condition in which the avatar and the shelf were tilted 45° in opposite directions, but this condition was not included in the analyses as we lacked the corresponding control condition where the avatar was upright and the shelf tilted 90° away from the vertical.

According to our working hypothesis, performance should be most affected in conditions engaging perspective-taking in a tilted orientation (i.e., roll rotation; avatar-tilt and both-tilt conditions). These conditions indeed correspond to situations where subjects need to mentally rotate themselves away from the gravity vector, so that the absence of a gravitational reference frame in 0g may influence, either negatively or positively, their performance. The shelf-tilt condition was critical in testing our hypothesis, since it served as a natural control for the avatar-tilt condition. Indeed, in these two conditions, the relative orientation of the avatar with respect to the shelf was identical. Consequently, should performance on the avatar-tilt condition—but not on the shelf-tilt condition—be affected in 0g, this would indicate that the impact on performance is not due to a trivial effect of visual incongruence between the shelf and the avatar orientation, but instead results specifically from the effects of microgravity on mental rotation processes.

### 2.4. Behavioral analysis

Preliminary analyses showed that response time distributions contained outliers. For each individual subject and condition, we systematically removed trials associated with response times that were 3 standard deviation above or below the mean. The mean and standard deviation were computed per subject and condition across all 3 sessions (preflight, inflight and postlight). Overall, less than 2% of the trials were identified as outliers and subsequently removed from the analyses.

For the extraction of 1g and 0g phases during the inflight session, we used acceleration parameters which were provided by Novespace (Merignac, France) and included gravity level (g-level) along the z-axis (floor-ceiling axis) at a millisecond resolution. The on-board time, which had been synchronized with the experiment computer CPU time, was used to attribute a g-level to every trial of the inflight session. By definition, a trial was considered to be performed in 0g if the g-level between the end of the instruction and the click release remained below 0.1g. With one parabola lasting about 20s, subjects typically completed 4 trials in 0g during each of these short periods. Trials performed during the periods of steady flight (after every 5 parabolas) were considered to be in 1g. For one subject, the main computer had to be restarted during one of the 1g periods. This subject completed the rest of their 1g trials after the very last parabola during steady flight.

### 2.5. Statistical analysis

We used Matlab 2017a (MathWorks, Natick, MA, USA) to perform all the statistical analyses. Data were gathered in a matrix containing, for each condition along the first dimension and each subject along the second dimension, the average response time or success rate computed over ~10 repetitions of the same condition. The number of repetitions per condition was slightly variable across subjects due to the outlier removal procedure and because subjects did not necessarily complete the same number of trials over the 15 parabolas. Given our relatively small sample size, we opted for non-parametric tests, i.e., Friedman test (equivalent of ANOVA with repeated measures) and Wilcoxon sign-rank tests for paired samples. All tests were two-tailed, unless specified otherwise. We considered p-values < 0.05 to be significant, unless post-hoc analyses with multiple comparisons were performed, in which case a Bonferroni correction was applied.

### 2.6. Rod-and-frame test

Subjects were seated in a complete dark room in front of a 17-cm computer display. A black cylinder placed between the subject and the screen formed an optical tunnel (distance subject-screen = 50 cm). One opening of the cylinder directly abutted the screen, while the other opening allowed subjects to rest their chin inside the tube. A black cloth covered the subject’s head to remove any peripheral visual cues. Subjects were required to keep their head upright during the test. The test itself consisted of a pre-programmed sequence of 10 rod-only trials followed by 10 rod-and-frame trials. On rod-only trials, the rod (luminescent bar, 4 cm long) was positioned pseudo-randomly at an angle of +40° or −40° from the vertical. Subjects were required to rotate the rod using the computer mouse until it aligned with the subjectively perceived vertical. Subjects used the mouse click to confirm their response. On rod-and-frame trials, the frame was tilted 18° in a clockwise (+18°) or counter-clockwise (−18°) direction from the vertical. The rod was positioned at an angle of either +20° or −20° from the vertical. To eliminate potential biases and learning effects, the combinations of rod/frame orientation were presented in a randomized order. The entire test lasted about three minutes and was preceded by a single familiarization trial.

We applied the well-established method developed by [44] to quantify each subject’s reliance on the visual field. According to this approach, performance at the rod-and-frame test can be reduced to 4 summary statistics. The constant error, μ, is the average (signed) deviation from the true vertical across all trials, regardless of frame or rod initial angle. The frame effect, φ, is the average deviation across all frame right-tilted trials (i.e., +18°), from which μ is subtracted. The rod starting position effect, ρ, is the average deviation across all rod initially right-tilted trials (i.e., +20°), from which μ is subtracted. Finally, the response consistency, σ, is the root-mean-square of the difference in deviation between same condition type, across all conditions. Following [44], we assumed that these followed a t-distribution, and assessed significance by computing 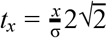, where *x* designates μ, φ or ρ. By definition, subjects who had φ significantly different from zero were considered to be field-dependent.

## 3. RESULTS

### 3.1. Performance on the ground

As an important initial step, we first analyzed performance on the ground to verify that the task engaged subjects cognitively. We used the data collected preflight (i.e., in normal gravity conditions) and assessed response time (RT) and success rate (SR) across experimental conditions. If the task requires subjects to perform own-body mental transformations with various degrees of difficulty, we should expect performance to vary between conditions. Accordingly, we found that the mean RT differed across condition type (non-parametric Friedman test with repeated measures, *χ*^2^(4)=19.85, p<0.001). In particular, we found that subjects were significantly slower when the avatar was tilted (avatar-tilt and both-tilt) compared to the no-tilt condition (Figure 2a; Wilcoxon sign-rank tests with Bonferroni correction and adjusted significance level of p<0.005). Subjects were also slower when the shelf alone was tilted (shelf-tilt) compared to no-tilt condition, indicating that the presence of allocentric cues influenced the mental transformation process. The success rate, defined as the percentage of correct responses in each condition, also varied systematically across the 5 conditions (Friedman test, *χ*^2^(4)=29.56, p<0.001). Follow-up pairwise comparisons revealed that subjects made significantly more errors when the avatar was tilted (avatar-tilt and both-tilt) compared to the no-tilt condition (Figure 2b; Wilcoxon sign-rank tests, p<0.005). These variations in performance both in terms of response time and success rate suggest that subjects engaged different cognitive processes across the 5 experimental conditions of the perspective-taking task.

**Figure 2.**
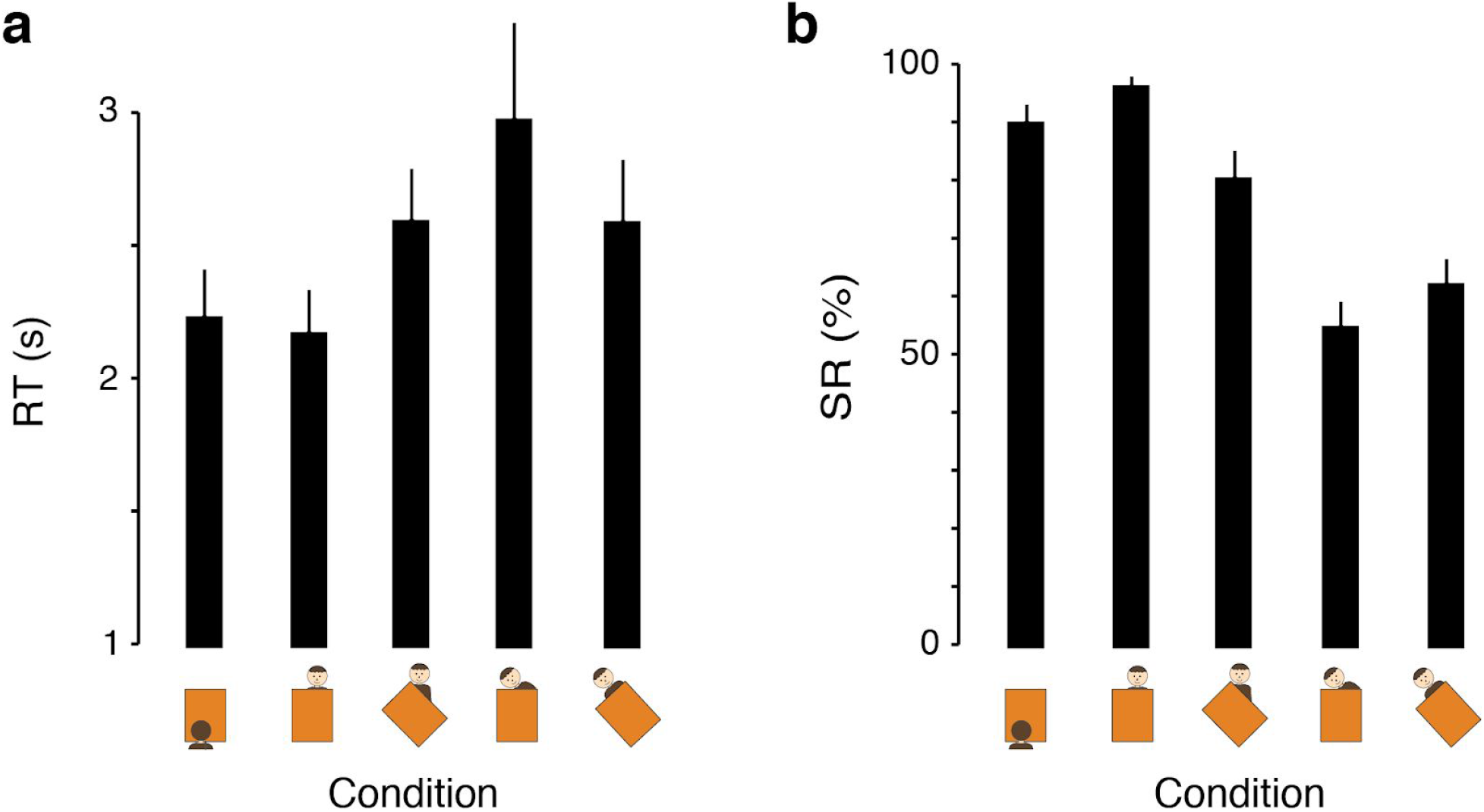
Performance on the ground. a) Mean response time (RT) in the 5 experimental conditions for the preflight session. RT is first averaged within subject, then combined across subjects to obtain a mean RT and the associated standard error (error bar). b) Same as a) for success rate (SR).

### 3.2. Effect of training: pre vs. postflight performance

Next we evaluated potential learning effects on subjects’ performance. Indeed, because data were collected over 3 separate sessions, performance may have improved due to repeated exposure to the task. We therefore compared performance pre and postflight. We found that RT was faster postflight for all conditions (Wilcoxon sign-rank tests, two-tailed, p<0.05 for all conditions, Figure 3a). By contrast, we did not find any systematic improvement in terms of success rate (Figure 3b). Except for an increase in SR postflight in the no-persp condition (sign-rank test, p=0.008), success rates associated with all other conditions were similar pre and postflight. The drop in RT between pre and postflight could be due to two main factors. One possibility is that subjects progressively improved at the mental transformations required in the task. This interpretation, however, is at odds with the observation that success rates were virtually unchanged in all conditions that required perspective-taking. The fact that RT decreased by roughly the same amount (0.54s on average) in all conditions, including the no-persp condition which did not involve any mental operation, also makes this explanation unlikely. A more likely account for the improved performance is an increased familiarity with the response apparatus. Anecdotal reports indeed indicated that subjects took some time to get used to manipulating the wireless mouse over the cardboard attached to their thigh (Figure 1a). If that is the case, we should expect the drop in RT to be independent of the condition type. Results indeed confirmed that the difference between pre and postflight mean RT did not vary significantly across conditions (Friedman test, *χ*^2^(4)=6.84, p=0.15). This strongly suggests that the likely source of faster performance postflight was a non-specific improvement in subjects’ ability to use the response apparatus.

**Figure 3.**
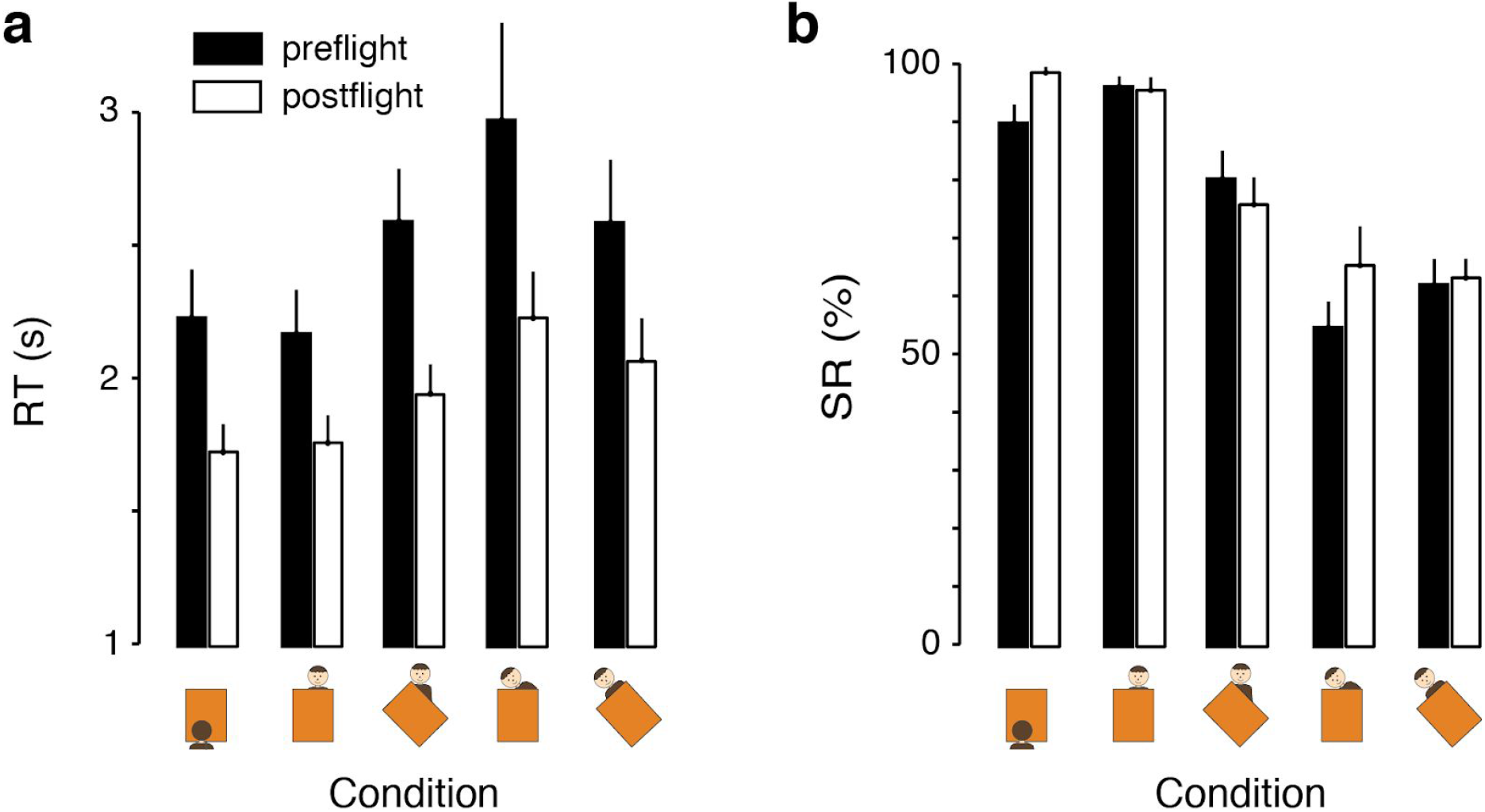
Performance pre and postflight. a) Mean response time (RT) in the 5 experimental conditions for pre (black) and postflight (white). b) Same as a) for success rate (SR).

### 3.3. Impact of inflight factors

Parabolic flights constitute a rather unusual setting for data collection: subjects are medicated (to alleviate motion sickness under altered gravity) and have to perform under stressful conditions, including high levels of noise and physical disturbances on the airplane. Although we had taken several measures to mitigate these nuisance factors (e.g., VR headset combined with over-the-ear headphones to isolate subjects from their environment), performance may still have been affected during the flight by parameters that were beyond our control. We evaluated whether subjects’ performance inflight during 1g periods differed from their baseline collected on the ground. Based on our previous analysis showing a learning effect between pre and postflight, we used postflight data as 1g baseline. Results revealed no difference in RT or SR between ground and 1g inflight performance (Wilcoxon sign-rank tests, p>0.05 for all conditions; Figure 4). This indicated that performance was not significantly impacted by inflight factors, and we therefore combined inflight 1g data with postflight data in the following analyses to gain statistical power.

**Figure 4.**
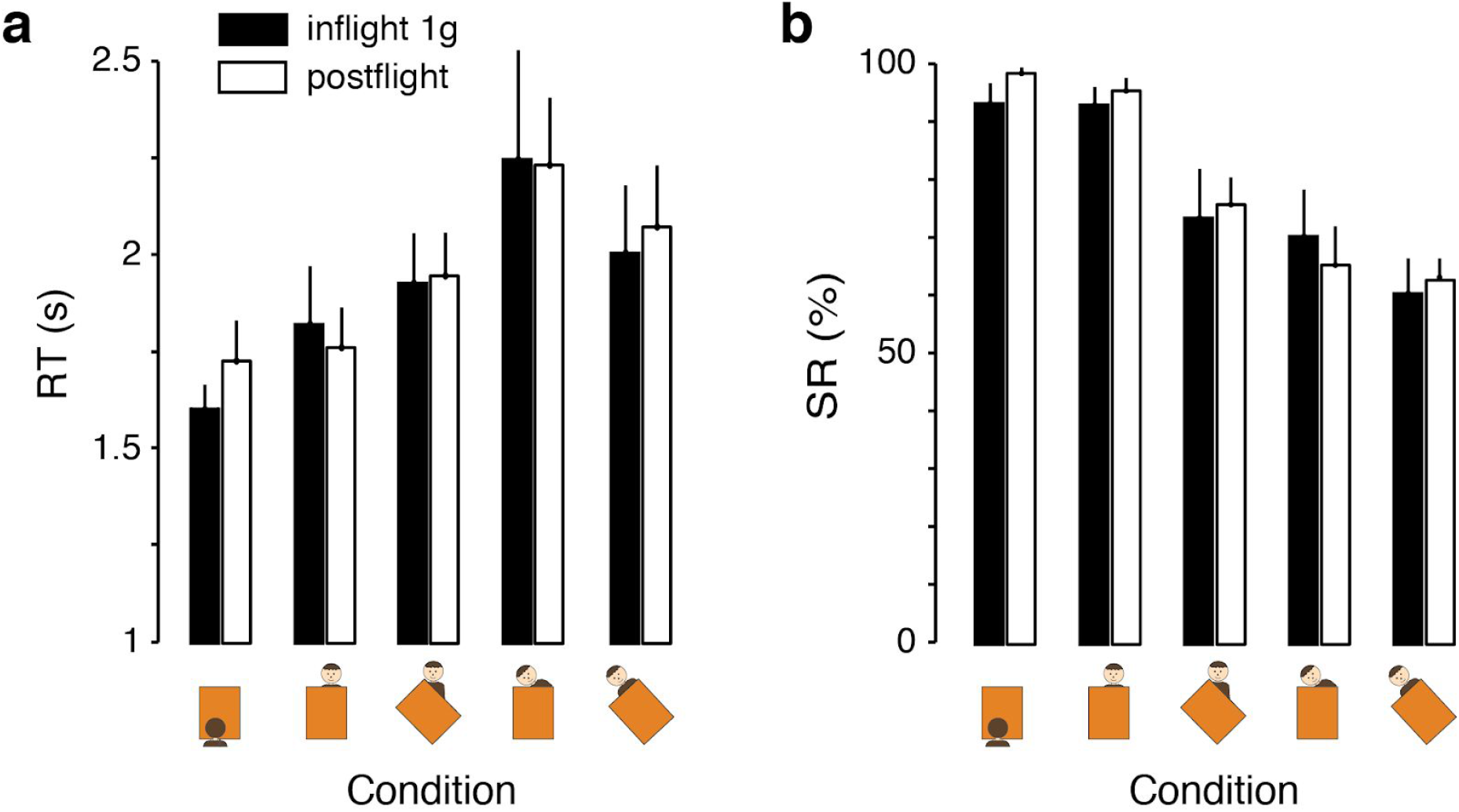
Performance inflight (1g) and postflight. a) Mean response time (RT) in the 5 experimental conditions for 1g inflight (black) and postflight (white). b) Same as a) for success rate (SR).

### 3.4. Effect of microgravity on performance

To test our main hypothesis of an effect of microgravity on perspective-taking abilities, we next compared 1g data to data collected in 0g during parabolas. We expected performance to be most influenced by weightlessness in tilted perspective-taking conditions, corresponding to situations where subjects mentally rotate their perspective away from the gravitational vector (roll rotations). Consistent with our prediction, we observed a specific decrease in RT in both the avatar-tilt and both-tilt conditions (difference in mean RT between 0g and 1g: 0.24 and 0.30 s, respectively; Figure 5a) but almost no difference in the no-persp, no-tilt and shelf-tilt conditions between 0g and 1g (difference in mean RT: −0.02, 0.02 and −0.02 s, respectively; Figure 5a). Statistical analyses corroborated these observations, as RT was significantly faster in 0g compared to 1g when grouping conditions that involved tilted perspective-taking in the frontal plane (avatar-tilt and both-tilt conditions; Wilcoxon sign-rank test, p=0.02). By contrast, RT associated with no or yaw-only rotation (grouping no-persp, no-tilt and shelf-tilt conditions) was not affected by microgravity (sign-rank test, p=0.46). In terms of SR, we did not find any improvement or degradation in performance between 0g and 1g for the grouped tilted perspective-taking conditions or the grouped control conditions (sign-rank tests, p=0.71 and p=0.90, respectively; Figure 5b). Together, these results indicate that subjects were faster, but as accurate in 0g compared to 1g in situations where they had to mentally project themselves in the frontal plane behind the shelf (yaw-roll rotation). The fact that this facilitation was not observed in control conditions requiring only a yaw rotation strongly suggest that the improved performance results from a specific effect of perceived gravity (or lack thereof in 0g) affecting roll rotation processes during perspective-taking.

**Figure 5.**
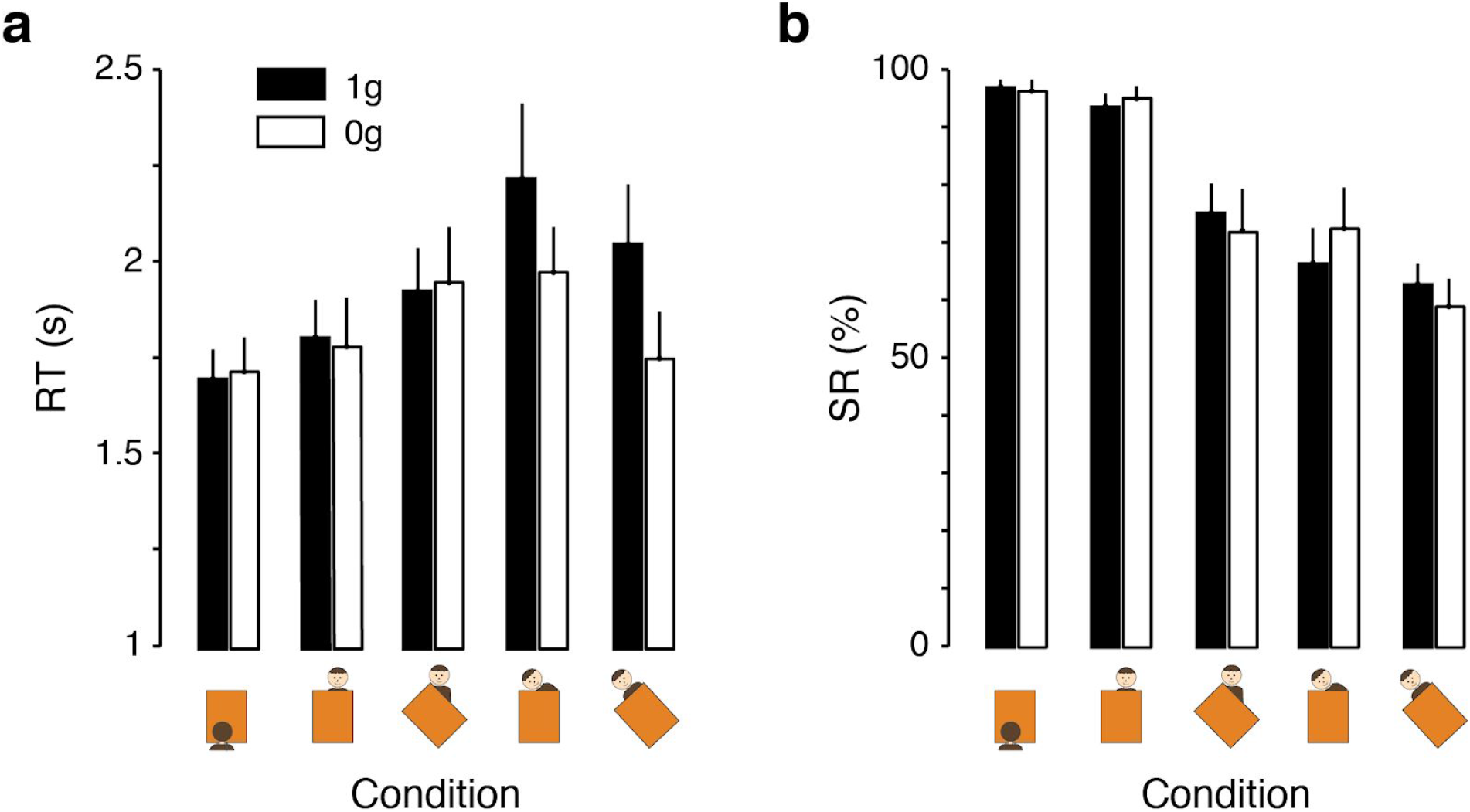
Performance in 1g versus 0g. a) Mean response time (RT) in the 5 experimental conditions for 1g (black) and 0g (white). b) Same as a) for success rate (SR). In both plots, 1g data combines postflight and inflight 1g.

### 3.5. Interaction between visual field-dependence and microgravity

Our experimental design included an independent measure of subjects’ dependence on the visual field. We reasoned that subjects who heavily rely on visual cues should be less, if at all impacted by weightlessness when solving the perspective-taking task. By contrast, we expected visuo-independent subjects, who presumably draw more heavily on vestibular cues to perform mental transformations, to be more affected by microgravity. We used the classic rod-and-frame test [43] to assess how much each individual subject was influenced by their visual field in perceiving the direction of gravity (Figure 6a). Subjects were classified as either field-dependent (FD) or non field-dependent (NFD) according to the metric developed by [44]; see Method). Based on this approach, we identified 4 (out of 11) subjects as being FD (Figure 6b). We analyzed FD and NFD subjects’ performance separately to test whether microgravity affected each group differently. Because sample sizes within both groups were small, we increased statistical power by grouping the two tilted perspective conditions before testing performance in 0g versus 1g. Consistent with what we had observed across all subjects, we found that NFD subjects were significantly faster at conditions involving a tilted perspective in 0g compared to 1g (Wilcoxon sign-rank test, p=0.01; Figure 6c). For FD subjects, however, we did not find any evidence of improved performance in terms of RT between 0g and 1g (sign-rank test, p=0.64; Figure 6c). As a control, we verified that both groups performed similarly in 0g and 1g in the grouped control conditions (sign-rank tests, p=0.23 for NFD, p=0.62 for FD). Taken together, these analyses provide evidence that the facilitation of egocentric transformation is selective to, or at the very least more pronounced in subjects that depend less on visual and more on vestibular cues.

**Figure 6.**
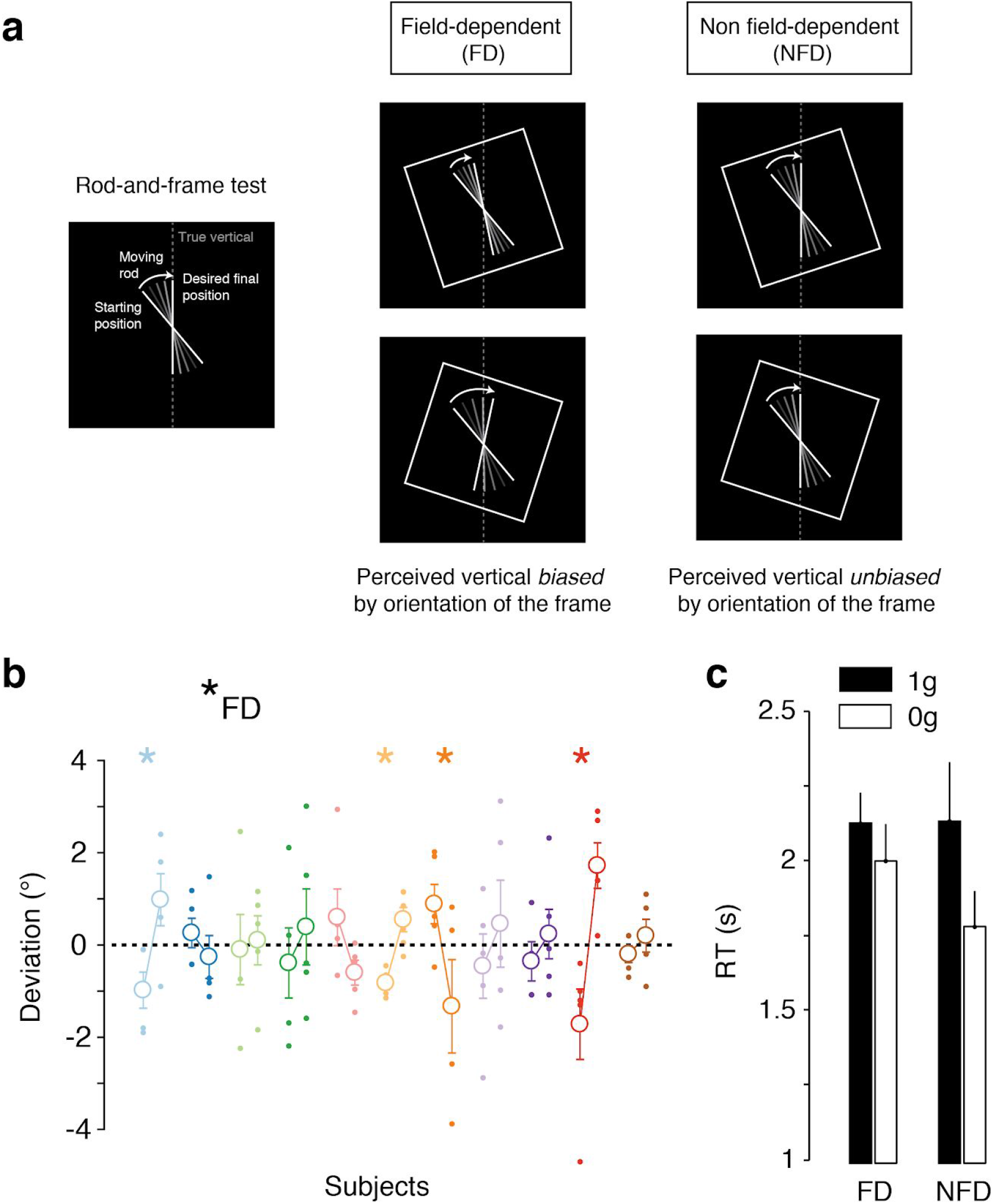
Performance in 1g versus 0g for field-dependent and non field-dependent subjects. a) Rod-and-frame test. Subjects are asked to rotate a moving rod from a random starting position until it is aligned with the perceived vertical (left). Subjects are classified as either field-dependent (FD, middle) or non field-dependent (NFD, right) depending on whether their responses are biased by the orientation of a frame surrounding the rod. b) Rod angular deviation on trials with frame. Each colored data pair corresponds to a single subject and shows the rod deviation for a frame tilted counterclockwise (left point) or clockwise (right). Each dot represents a single trial; open circles show the mean and standard error. The average deviation across all trials per subject has been removed to get a common baseline. Stars designate subjects identified as FD per Nyborg and Isaksen’s method. c) Mean response time (RT) for tilted-perspective conditions (combining avatar-tilt and both-tilt) for 1g (black) and 0g (white) plotted separately for FD and NFD subjects.

## 4. DISCUSSION

The effect of microgravity on spatial cognition is a long standing question in space neuroscience [4, 45–47]. Not only does this question matter for operational purposes, it is also key for understanding the role of gravity in fundamental cognitive processes such as mental imagery [6, 48]. Many studies in the past have attempted to characterize 0g performance in mental rotation tasks [16]. Results, however, remain highly equivocal, with some studies finding a facilitation in microgravity [21–23], while others concluded in no effect [28–30], or even a decline in performance [16, 31]. Moreover, the question of mental imagery in microgravity has almost exclusively been asked from a theoretical standpoint, employing relatively simplistic stimuli [31, 41, 42]. The goal of this study was therefore two-fold. First, we aimed to resolve the debate regarding the negative or positive impact of weightlessness specifically on perspective-taking [19, 49]. Second, we sought to characterize performance in an operationally-relevant setting much more likely to translate to real-world situations faced by astronauts.

Using a novel paradigm leveraging virtual reality aboard a parabolic flight, we found that weightlessness has a facilitatory effect on perspective-taking performance (Figure 5) that is selective to rotations in the frontal plane (i.e., roll rotation around subjects’ line of sight). Conditions that solely engaged rotations around the vertical axis (yaw rotation), on the other hand, were unaffected in 0g. It is worth mentioning that a similar effect has been previously reported in cosmonauts aboard the MIR station on a simpler mental object-rotation task [21–23]. Results of this study, however, were obscured by large individual differences within the cohort as well as significant learning effects throughout the experiment. Although we did observe learning in our task (Figure 3), we controlled for this effect in two ways: (1) inflight 1g and 0g data were collected in a single experimental session and over rapidly alternating periods, thus minimizing the extent of learning [33], and (2) postflight data collected on the ground were compared to inflight 1g data to ensure that performance was similar during and after the flight. Regardless, even if learning had occurred during the flight, this could not explain the facilitation observed in 0g, since the collection of inflight 0g data always alternated with inflight 1g data collection.

Why are mental rotations in the frontal plane selectively facilitated in 0g? One interpretation that has been put forth [21–23] relates to the idea of motoric embodiment [24, 50]. According to this framework, physical constraints including gravitational effects directly impact internal processes engaged in operations such as mental imagery [16]. Put simply, if a movement is physically impossible or difficult to perform in reality due to mechanical or other environmental constraints, then the process of imagining that same movement is equally laborious. For instance, when subjects are asked to mentally picture their hand rotating in directions not physically achievable, response times tend to differ greatly from more common movements [51, 52]. Along the same lines, subjects asked to take the perspective of an avatar respond faster when their own body is being rotated in the same direction as the mental rotation needed to achieve the desired perspective [53].

These findings underscore the tight relationship between imagined and real actions [26]. In the case of microgravity, subjects evolve in an environment where they are free to float around the cabin. This releases the physical constraints on how their body can move in space, and rotations especially around an individual’s front-back axis become “unsurprising”. This effect may explain why in the present experiment response times were faster only on conditions requiring participants to mentally rotate themselves in the frontal plane. Despite the fact that participants were seated and loosely attached (hence unable to completely free-float), the mere absence of otolithic cues in 0g may have been sufficient to trigger facilitatory effects on mental rotations [16]. Indeed, previous studies have reported that astronauts perform similarly on a mental representation task while free-floating or while seated and attached [54].

Alternatively, decreased reaction times in 0g might result from other less specific effects. In particular, sensorimotor changes in 0g have been repeatedly reported. For instance, drawings were performed faster in 0g compared to 1g [55]. If sensorimotor facilitation had occurred in the present experiment, reaction times in all test conditions, including the control conditions, should have been observed, but this was not the case. Further, static visual cues have been shown to affect embodiment and hence performance in an egocentric mental transformation task [38]. However, if this were also the case in the present study, a similar facilitation should have been observed in the shelf-tilt condition designed to match the relative orientation of the shelf with respect to the avatar. In addition, the fact that 0g impacted only a subset of experimental conditions in our task constitutes compelling evidence that the observed effects are not trivially explained by stress or other nuisance factors present during the parabolic flight [32, 56]. In fact, it has been shown that when participants are exposed to an acute laboratory stress while performing a virtual reality perspective-taking task, they respond on the contrary more slowly [32]. It is worth pointing out that, in the present study, we did not find any evidence of flight-induced changes in performance when comparing ground to inflight 1g data (Figure 4). We attribute this observation to the highly immersive nature of the task which successfully isolated subjects from cabin disturbances.

Another insight gleaned from our results is the interaction between the effects of microgravity and visual reliance. As stated by [6]: “there are two general categories of individuals: those who come to rely on the visual verticals of the spacecraft to gauge their body orientation [field-dependent] and those who rely on the location of their feet for the direction of down [non field-dependent]. The former feel upright when they are aligned with the architectural verticals of the spacecraft with their head toward the ‘ceiling’, the latter feel upright regardless of their orientation to the spacecraft.” Consistent with this qualitative dissociation, we indeed found that the facilitation of egocentric transformations in our task was more pronounced in non field-dependent subjects (Figure 6), as determined by an independent Rod-and-Frame test [43]. These subjects likely rely more heavily on vestibular and haptic/proprioceptive cues compared to visual cues to assess verticality [1, 57]. It is therefore not entirely surprising that weightlessness, associated with the unloading of the vestibular system and the partial loss of haptic/proprioceptive feedback, would particularly influence their performance at the task.

What is the neural basis of the facilitation of egocentric mental rotation in microgravity? On Earth, spatial representations are deeply anchored to gravity via a multisensory process that integrates vestibular, proprioceptive and visual signals altogether [2]. Manipulating objects and reference frames [35], as in the case of mental imagery, is therefore influenced by these interacting variables [58]. The involvement of the vestibular system in spatial cognition is particularly well-established [48]. For instance, vestibular stimulation is known to facilitate spatial judgments, especially in the case of own-body rotations [37, 53]. The parietal cortex may also play a critical role in the observed effects given its major contribution to multisensory spatial processing [59, 60]. Furthermore, gravity has been shown to modulate the neural representation of object tilt in the parietal cortex [61], which may have solicited in the tilted perspective-taking conditions. However, the influence of microgravity on mental imagery likely involves a distributed network of brain regions, and remains to be fully characterized.

Regardless of its neural correlate, the selective improvement of performance we observe in 0g are likely to have direct implications for space operations. To our knowledge, no study has attempted to characterize the influence of microgravity on mental imagery in a realistic, collaborative setting. Virtual reality, which has previously been employed to investigate navigation and orientation abilities inside simulated spacecrafts [62, 63], provides a rich yet highly controlled environment to recreate operational conditions close to real-life situations encountered by astronauts. This allowed us to assess for the first time the potential impact of weightlessness on crew interactions during a complex cooperative task. Our data suggest that perspective-taking abilities engaged during such interactions would not be negatively impacted by microgravity. If the present results were further confirmed, they would alleviate some of the human factors concerns regarding long-duration space missions, which will increasingly rely on the successful coordination of complicated tasks by large groups of agents.

## Acknowledgments

This project was supported by the French space agency (Centre National d’Etudes Spatiales, CNES, France). We thank Sébastien Rouquette and Maurice Marnat for providing most of the hardware (laptops and virtual reality equipment). We also thank Sébastien Barde and Alain Maillet for constructive discussions in the early stages of the project, as well as Dr. Anne Pavy-Le Traon and Guillemette Gauquelin-Koch for administrative and financial aspects. Finally, we thank Novespace (Mérignac, France) responsible for the parabolic flight campaign, and in particular Thomas Villatte for his technical guidance throughout the project.

## REFERENCES

1. Friederici AD, Levelt WJ. Spatial reference in weightlessness: perceptual factors and mental representations. Percept Psychophys. 1990;47: 253–266.

2. Berthoz A, Viaud-Delmon I. Multisensory integration in spatial orientation. Curr Opin Neurobiol. 1999;9: 708–712.

3. Harris LR, Herpers R, Hofhammer T, Jenkin M. How much gravity is needed to establish the perceptual upright? PLoS One. 2014;9: e106207.

4. Clément G, Reschke MF. Neuroscience in Space. Springer Science & Business Media; 2010.

5. Bock O. Problems of sensorimotor coordination in weightlessness. Brain Res Brain Res Rev. 1998;28: 155–160.

6. Lackner JR, DiZio P. Human orientation and movement control in weightless and artificial gravity environments. Exp Brain Res. 2000;130: 2–26.

7. Bock O, Fowler B, Comfort D. Human sensorimotor coordination during spaceflight: an analysis of pointing and tracking responses during the “Neurolab” Space Shuttle mission. Aviat Space Environ Med. europepmc.org; 2001;72: 877–883.

8. Semjen A, Leone G, Lipshits M. Motor timing under microgravity. Acta Astronaut. Elsevier; 1998;42: 303–321.

9. De la Torre GG. Cognitive neuroscience in space. Life. 2014;4: 281–294.

10. Pattyn N, Migeotte P-F, Demaeseleer W, Kolinsky R, Morais J, Zizi M. Investigating human cognitive performance during spaceflight. adsabs.harvard.edu; 2005.

11. Clément G, Lathan C, Lockerd A. Perception of depth in microgravity during parabolic flight. Acta Astronaut. 2008;63: 828–832.

12. Clément G, Demel M. Perceptual reversal of bi-stable figures in microgravity and hypergravity during parabolic flight. Neurosci Lett. 2012;507: 143–146.

13. Leone G. The effect of gravity on human recognition of disoriented objects. Brain Res Brain Res Rev. 1998;28: 203–214.

14. Papaxanthis C, Pozzo T, Kasprinski R, Berthoz A. Comparison of actual and imagined execution of whole-body movements after a long exposure to microgravity. Neurosci Lett. 2003;339: 41–44.

15. Pozzo T, Papaxanthis C, Stapley P, Berthoz A. The sensorimotor and cognitive integration of gravity. Brain Res Brain Res Rev. 1998;28: 92–101.

16. Grabherr L, Mast FW. Effects of microgravity on cognition: The case of mental imagery. J Vestib Res. 2010;20: 53–60.

17. De La Torre GG, van Baarsen B, Ferlazzo F, Kanas N, Weiss K, Schneider S, et al. Future perspectives on space psychology: Recommendations on psychosocial and neurobehavioural aspects of human spaceflight. Acta Astronaut. Elsevier; 2012;81: 587–599.

18. Tversky B, Hard BM. Embodied and disembodied cognition: spatial perspective-taking. Cognition. 2009;110: 124–129.

19. Dumontheil I, Küster O, Apperly IA, Blakemore S-J. Taking perspective into account in a communicative task. Neuroimage. 2010;52: 1574–1583.

20. Menchaca-Brandan MA, Liu AM, Oman CM, Natapoff A. Influence of Perspective-taking and Mental Rotation Abilities in Space Teleoperation. Proceedings of the ACM/IEEE International Conference on Human-robot Interaction. New York, NY, USA: ACM; 2007. pp. 271–278.

21. Clement G, Berthoz A, Lestienne F. Adaptive changes in perception of body orientation and mental image rotation in microgravity. Aviat Space Environ Med. 1987;58: A159–63.

22. Matsakis Y, Lipshits M, Gurfinkel V, Berthoz A. Effects of prolonged weightlessness on mental rotation of three-dimensional objects. Exp Brain Res. 1993;94: 152–162.

23. Harm DL, Parker DE. Perceived self-orientation and self-motion in microgravity, after landing and during preflight adaptation training. J Vestib Res. 1993;3: 297–305.

24. Amorim M-A, Isableu B, Jarraya M. Embodied spatial transformations: “body analogy” for the mental rotation of objects. J Exp Psychol Gen. 2006;135: 327–347.

25. Mast FW, Bamert L, Newby N. Mind over Matter? Imagined Body Movements and Their Neuronal Correlates. In: Mast F, Jäncke L, editors. Spatial Processing in Navigation, Imagery and Perception. Boston, MA: Springer US; 2007. pp. 353–368.

26. Jeannerod M. Neural simulation of action: a unifying mechanism for motor cognition. Neuroimage. 2001;14: S103–9.

27. Dyde RT, Jenkin MR, Jenkin HL, Zacher JE, Harris LR. The effect of altered gravity states on the perception of orientation. Exp Brain Res. 2009;194: 647–660.

28. Leone G, Lipshits M, Gurfinkel V, Berthoz A. Is there an effect of weightlessness on mental rotation of three-dimensional objects? Brain Res Cogn Brain Res. 1995;2: 255–267.

29. Dalecki M, Dern S, Steinberg F. Mental rotation of a letter, hand and complex scene in microgravity. Neurosci Lett. 2013;533: 55–59.

30. Gurfinkel VS, Lestienne F, Levik YS, Popov KE, Lefort L. Egocentric references and human spatial orientation in microgravity. Exp Brain Res. 1993;95: 343–348.

31. Grabherr L, Karmali F, Bach S, Indermaur K, Metzler S, Mast FW. Mental own-body and body-part transformations in microgravity. J Vestib Res. 2007;17: 279–287.

32. Richardson AE, VanderKaay Tomasulo MM. Influence of acute stress on spatial tasks in humans. Physiol Behav. 2011;103: 459–466.

33. Wollseiffen P, Vogt T, Abeln V, Strüder HK, Askew CD, Schneider S. Neuro-cognitive performance is enhanced during short periods of microgravity. Physiol Behav. 2016;155: 9–16.

34. Zacks JM, Mires J, Tversky B, Hazeltine E. Mental Spatial Transformations of Objects and Perspective. Spat Cogn Comput. Hingham, MA, USA: Kluwer Academic Publishers; 2001;2: 315–332.

35. Klatzky RL. Allocentric and Egocentric Spatial Representations: Definitions, Distinctions, and Interconnections. In: Freksa C, Habel C, Wender KF, editors. Spatial Cognition: An Interdisciplinary Approach to Representing and Processing Spatial Knowledge. Berlin, Heidelberg: Springer Berlin Heidelberg; 1998. pp. 1–17.

36. Grabherr L, Cuffel C, Guyot J-P, Mast FW. Mental transformation abilities in patients with unilateral and bilateral vestibular loss. Exp Brain Res. 2011;209: 205–214.

37. Falconer CJ, Mast FW. Balancing the mind: vestibular induced facilitation of egocentric mental transformations. Exp Psychol. 2012;59: 332–339.

38. Preuss N, Harris LR, Mast FW. Allocentric visual cues influence mental transformation of bodies. J Vis. 2013;13: 14.

39. Witkin HA, Goodenough DR. Cognitive styles: essence and origins. Field dependence and field independence. Psychol Issues. 1981; 1–141.

40. Shepard RN, Metzler J. Mental rotation of three-dimensional objects. Science. 1971; 171: 701–703.

41. Eddy DR, Schiflett SG, Schlegel RE, Shehab RL. Cognitive performance aboard the life and microgravity spacelab. Acta Astronaut. 1998;43: 193–210.

42. de Schonen S, Leone G, Lipshits M. The face inversion effect in microgravity: is gravity used as a spatial reference for complex object recognition? Acta Astronaut. 1998;42: 287–301.

43. Witkin HA, Asch SE. Studies in space orientation; further experiments on perception of the upright with displaced visual fields. J Exp Psychol. 1948;38: 762–782.

44. Nyborg H, Isaksen BO. A method for analysing performance in the rod-and-frame test. II Test of the Statistical Model. Scand J Psychol. 1974;15: 124–126.

45. Fowler B, Comfort D, Bock O. A review of cognitive and perceptual-motor performance in space. Aviat Space Environ Med. 2000;71: A66–8.

46. Casler JG, Cook JR. Cognitive performance in space and analogous environments. Int J Cogn Ergon. 1999;3: 351–372.

47. Harris LR, Jenkin M, Jenkin H, Zacher JE, Dyde RT. The effect of long-term exposure to microgravity on the perception of upright. NPJ Microgravity. 2017;3: 3.

48. Bigelow RT, Agrawal Y. Vestibular involvement in cognition: Visuospatial ability, attention, executive function, and memory. J Vestib Res. 2015;25: 73–89.

49. Keysar B, Barr DJ, Balin JA, Brauner JS. Taking perspective in conversation: the role of mutual knowledge in comprehension. Psychol Sci. 2000;11: 32–38.

50. Wexler M, Kosslyn SM, Berthoz A. Motor processes in mental rotation. Cognition. 1998;68: 77–94.

51. Tao W, Liu Q, Huang X, Tao X, Yan J, Teeter CJ, et al. Effect of degree and direction of rotation in egocentric mental rotation of hand: an event-related potential study. Neuroreport. 2009;20: 180–185.

52. Parsons LM. Temporal and kinematic properties of motor behavior reflected in mentally simulated action. J Exp Psychol Hum Percept Perform. 1994;20: 709–730.

53. Deroualle D, Borel L, Devèze A, Lopez C. Changing perspective: The role of vestibular signals. Neuropsychologia. 2015;79: 175–185.

54. Lathan C, Wang Z, Clément G. Changes in the vertical size of a three-dimensional object drawn in weightlessness by astronauts. Neurosci Lett. 2000;295: 37–40.

55. Clément G, Lathan C, Lockerd A, Bukley A. Mental representation of spatial cues in microgravity: Writing and drawing tests. Acta Astronaut. 2009;64: 678–681.

56. Cane JE, Ferguson HJ, Apperly IA. Using perspective to resolve reference: The impact of cognitive load and motivation. J Exp Psychol Learn Mem Cogn. 2017;43: 591–610.

57. Luyat M, Mobarek S, Leconte C, Gentaz E. The plasticity of gravitational reference frame and the subjective vertical: peripheral visual information affects the oblique effect. Neurosci Lett. 2005;385: 215–219.

58. Lackner JR, DiZio P. Vestibular, proprioceptive, and haptic contributions to spatial orientation. Annu Rev Psychol. 2005;56: 115–147.

59. Chang SWC, Snyder LH. Idiosyncratic and systematic aspects of spatial representations in the macaque parietal cortex. Proc Natl Acad Sci U S A. 2010;107: 7951–7956.

60. Seilheimer RL, Rosenberg A, Angelaki DE. Models and processes of multisensory cue combination. Curr Opin Neurobiol. 2014;25: 38–46.

61. Rosenberg A, Angelaki DE. Gravity influences the visual representation of object tilt in parietal cortex. J Neurosci. 2014;34: 14170–14180.

62. Aoki H, Ohno R, Yamaguchi T. The effect of the configuration and the interior design of a virtual weightless space station on human spatial orientation. Acta Astronaut. 2005;56: 1005–1016.

63. Oman C. Spatial Orientation and Navigation in Microgravity. In: Mast F, Jäncke L, editors. Spatial Processing in Navigation, Imagery and Perception. Boston, MA: Springer US; 2007. pp. 209–247.

65. McIntyre J, Zago M, Berthoz A, Lacquaniti F. Does the brain model Newton’s laws? Nat Neurosci. 2001;4: 693–694.

